# Effects of treatment with enrofloxacin or tulathromycin on fecal microbiota composition and function of dairy calves

**DOI:** 10.1101/688788

**Authors:** Carla Foditsch, Richard V.V. Pereira, Julie D. Siler, Craig Altier, Lorin D. Warnick

**Author notes:** Corresponding author: Carla Foditsch, Department of Population Medicine and Diagnostic Sciences, College of Veterinary Medicine, Cornell University, Ithaca, NY 14853-6401.

## Abstract

The increasing concerns with antimicrobial resistance highlights the need for studies evaluating the impacts of antimicrobial use in livestock on antimicrobial resistance using new sequencing technologies. Through shotgun sequencing, we investigated the changes in the fecal microbiome composition and function, with a focus on functions related to antimicrobial resistance, of dairy calves. Heifers 2 to 3 weeks old, which were not treated with antibiotics by the farm before enrollment, were randomly allocated to one of three study groups: control (no treatment), enrofloxacin, or tulathromycin. Fecal samples were collected at days 4, 14, 56 and 112 days after enrollment, and DNA extraction and sample preparation and sequencing was conducted. The effect of antibiotic treatment on each taxon and functional level by time (including Day 0 as a covariate) revealed few changes in the microbiota. At the genus level, enrofloxacin group had higher abundance of *Blautia, Coprococcus* and *Desulfovibrio* and lower abundance of *Bacteroides* when compared to other treatment groups. The SEED database was used for functional analyses, which showed that calves in the enrofloxacin group started with a higher abundance of “Resistance to antibiotics and toxic compounds” function on Day 0, however an increase in antibiotic resistance genes after treatment with enrofloxacin was not observed. “Resistance to Fluoroquinolones” and “Erythromycin resistance”, of relevance given the study groups, were not statistically different in abundance between treatment groups. “Resistance to fluoroquinolones” increased during the study period regardless of treatment group. Despite small differences over the first weeks between treatment groups, at Day 112 the microbiota composition and functional profile was similar among all study groups. These findings show that metaphylaxis treatment of dairy calves with either enrofloxacin or tulathromycin have minimal impacts on the microbial composition and functional microbiota of dairy calves over time.

## Introduction

There is urgent need for the judicious use of antimicrobial drugs to extend the effectiveness of currently available therapies [1,2]. Antimicrobial resistance (AMR) is a natural phenomenon, and resistance genes are present even in places that were never inhabited by humans [3,4] and in wild animals [5]. However, the use of antibiotics in human and animal medicine, as well as in plant agriculture and animal production systems, has resulted in an acceleration in the selection and spread of antimicrobial resistance [1,6]. Antimicrobial resistant enteric bacteria can be transmitted from livestock to humans through the fecal-oral route, or contamination in the food chain and environment [1,7,8]. Together with prevention of disease and infectious agent transmission, the prudent use of antibiotics is extremely important to preserve treatment effectiveness and decrease the dissemination of resistant bacteria.

Antimicrobials may be used to control and prevent the spread of the disease in production animals at high risk of developing a bacterial infectious disease. This practice is referred to as metaphylaxis. In the U.S., several drugs are approved for this use in cattle, including in dairy calves, at risk of bovine respiratory disease (BRD). BRD is known to be a common cause of disease in dairy calves at 2-3 weeks of age [9,10]. In 2014 in the United States 12% of pre-weaned calves were affected with respiratory disease and almost 95% of those were treated with antimicrobial drugs [11]. Macrolides and florfenicol were the drugs of choice on 18.2 and 15.1 percent of the farms to treat BRD, followed by penicillin (8.1%) and fluoroquinolones (6.6 %) [11]. In our study, we focused on enrofloxacin and tulathromycin, a fluoroquinolone and a macrolide, respectively. These are injectable, single dose antimicrobials, labeled for treatment and control of BRD.

Fluoroquinolones are broad-spectrum antimicrobial drugs that target essential bacterial enzymes DNA gyrase and DNA topoisomerase IV. Mutations in these bacterial enzymes can confer resistance, as can plasmids carrying resistance genes that protect the cells from the bactericidal effects of quinolones [12,13]. Fluoroquinolones were developed about 40 years ago and their use increased worldwide in the 1990s for treatment of Gram-negative infections in humans [14,15]. Examples of FDA-approved fluoroquinolones for use in human medicine are ciprofloxacin and levofloxacin. In 1988, the FDA approved the veterinary use of a fluoroquinolone, enrofloxacin, for dogs and cats [16]. Later, in 1998, it was approved for treating bovine respiratory disease in cattle. The use of enrofloxacin in poultry was banned in 2005 and in 2008 its use in female dairy cattle was restricted to nonlactating animals up to 20 months old. Extra-label use of enrofloxacin is strictly prohibited. Enrofloxacin has also been approved for treatment and control of swine respiratory disease since 2008. [16].

Macrolides are mainly bacteriostatic. They inhibit bacterial protein synthesis and the spectrum of action of drugs within this class varies. In human medicine, erythromycin has been available since 1952, and other current drugs in the same class are azithromycin, clarithromycin and telithromycin [17]. In veterinary medicine, erythromycin and tylosin are used to treat companion animals, ruminants, swine, and poultry [16]. Tulathromycin, the macrolide used in our study, was approved by the FDA in 2005 to treat and control respiratory disease in cattle and swine [16].

Antimicrobial treatment is important for treatment and prevention of infections by pathogenic bacteria, but may also select for resistant strains and increase the prevalence of resistance genes that may be transferred to other bacterial strains [1,18]. The majority of antimicrobial resistance studies have focused on the effect of antibiotic treatment on individual bacterial strains [19,20]. Next Generation Sequencing is a relatively novel technique that is helping expand our knowledge of the impact of antibiotic treatment on the gut microbiome and the prevalence of bacterial genes in diverse bacterial populations [5,21,22]. There is limited published information on the impacts of antimicrobial drug treatment on microbial function in dairy calves, leaving a knowledge gap that limits the thorough evaluation of the impact of the use of wide spectrum antimicrobial drugs in young calves.

The objective of this study was to longitudinally characterize the effect of enrofloxacin or tulathromycin metaphylactic treatment on the fecal microbiome and microbial function of preweaned dairy calves, focusing on functions related to antimicrobial resistance.

## Material and Methods

### Ethics statement

The research protocol was reviewed and approved by the Institutional Animal Care and Use Committee of Cornell University (Protocol number: 2014-0094). The drug administration to calves housed on the commercial dairy farm and the collection of fecal samples was authorized by the farm owner, who was aware of all experimental procedures. The herd veterinarian determined the need for metaphylactic treatment and the drugs were administered according to label directions.

### Farm management

The study was conducted from May to November 2015 at a commercial dairy farm that milked 1,200 Holstein cows near Ithaca, New York, USA. The farm was selected because preventive antimicrobial treatment was indicated based on a prior history of calfhood respiratory disease identified by the herd veterinarian. Routine calf management was performed by farm employees. Newborn calves received four quarts of colostrum within the first hours of life and were bottle fed pasteurized non-saleable milk twice a day until weaning at approximately 56 days of age. Water was offered ad libitum in buckets. Heifer calves were kept in individual hutches or pens with sawdust bedding during the preweaning period (first 8 weeks of life) and then moved to group pens. Health-related events (e.g. otitis, pneumonia and diarrhea) were treated as needed by farm employees and recorded in the farm’s herd management software (Dairy-Comp 305; Valley Ag Software, Tulare, CA, USA). The research group obtained these health-related and treatment records from the herd management software and from the farm’s drug use notebook to increase data accuracy.

A randomized field trial study design was used. Calves 2 to 3 weeks old, which had not been treated with antimicrobials before enrollment, were randomly allocated within cohort (enrollment day) to one of three study groups: control (**CON**, no antimicrobial), enrofloxacin (**ENR**, Baytril 100, Bayer Corp. Agricultural Division, Shawnee Mission, KS, USA) or tulathromycin (**TUL**, Draxxin, Zoetis, Floham Park, NJ, USA). A total of 84 heifers were enrolled in the trial: 26 allocated to CON, 28 to ENR and 27 to TUL, distributed in 6 cohorts (6 enrollment days). Fecal samples were collected directly from the rectum of individual calves starting before the administration of the antimicrobial treatment on enrollment day (Day 0) and on 4, 14, 56, and 112 days after enrollment. Fecal samples were transported in a cooler with ice packs to the laboratory at the Cornell campus in Ithaca, NY, where they were aliquoted and stored at -20°C until DNA extraction. For whole genome sequencing, a subset of 12 calves per study group was randomly selected after excluding calves with missing samples or incomplete data, and calves treated with antimicrobials or other drugs by farm personnel or the herd veterinarian after enrollment.

### DNA extraction

Fecal samples were thawed and total DNA was extracted using the MoBio PowerSoil DNA isolation kit (MoBio Laboratories, Carlsbad, CA, USA) following the protocol previously used by Pereira *et al*. with few modifications [23]. Briefly, approximately 50 mg of feces was transferred to the PowerBead Tube, which was heat treated at 65°C for 10 minutes and then 95°C for 10 minutes. PowerBead Tubes were vortexed horizontally using the MoBio Vortex Adapter tube holder at a maximum speed for 10 minutes. The remaining DNA extraction procedure followed the standard protocol supplied by the kit manufacturer. The DNA concentration and purity were evaluated by optical density using a NanoDrop ND-1000 spectrophotometer (NanoDrop Technologies, Rockland, DE, USA) at wavelengths of 230, 260 and 280 nm. The eluted DNA was stored at -20°C until further processing.

### Library Preparation and Illumina HiSeq Sequencing

To increase measurement accuracy for the concentration for the DNA library, the starting DNA library was quantified using a fluorometric-based method, the Qubit dsDNA HS Assay system (Life Technologies Corporation, Carlsbad, CA, USA). An aliquot of each DNA sample was diluted to 0.2 ng/μl and used as input DNA to the Nextera XT DNA Library Preparation Kit (Illumina Inc. San Diego, CA), according to manufacturer’s recommendations. As a quality control step after PCR-cleanup, a subset of libraries was run on an Agilent Technology Bioanalyzer to check the size distribution (bp). After library normalization and pooling, sequencing was performed at the Genomics Facility of the Cornell University Institute of Biotechnology with an Illumina HiSeq2500 platform, operating in Rapid Run Mode, paired-end 2 × 250 bp, one sample pool in both lanes of a two-lane flowcell.

### MG-RAST Analysis

The paired-end sample sequences were obtained from the Cornell University Institute of Biotechnology as two individual fastq files with one file per end. The two files were concatenated using Python and uploaded to the Metagenomics Rapid Annotation using Subsystem Technology (MG-RAST), which is an open-source server that analyzes sequencing data and provides taxonomic and functional classification [24]. In summary, before submission for analyses at MG-RAST, standard screening and quality control procedures were selected, including removal of artificial replicate sequences, host (*Bos Taurus*) sequences and low quality sequences. The Refseq and SEED Subsystems databases were selected for taxonomic distributions (Phylum, Class, Order, Family, Genus and Species) and functional assignments (4 levels of hierarchy), respectively, as outlined in Figure 1.

**Fig 1.**
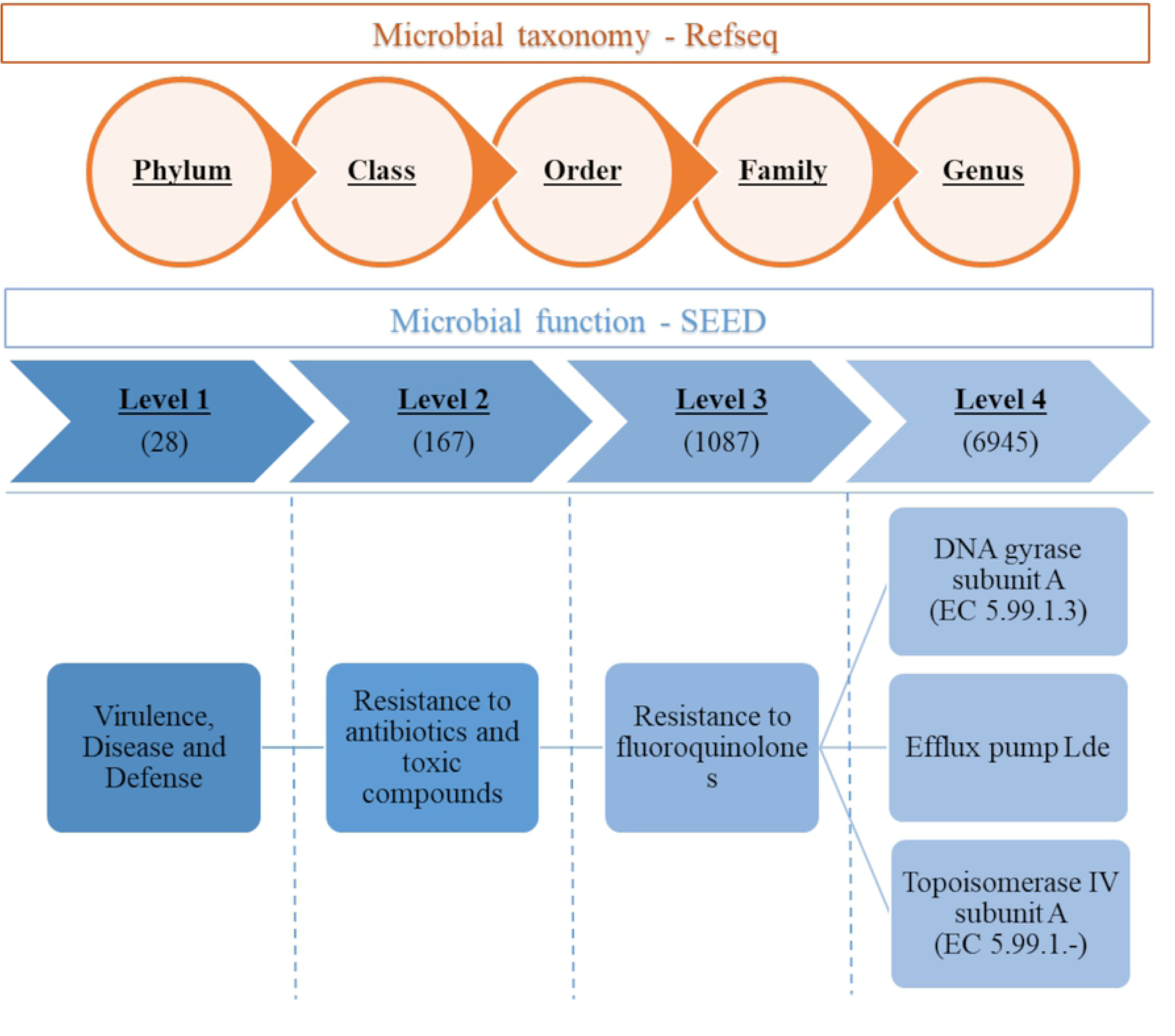
Scheme of the MG-RAST analysis, showing the levels of hierarchy. Refseq database was selected for taxonomic distributions (Phylum, Class, Order, Family, and Genus) and SEED Subsystems database was used for the functional analysis (Levels 1 to 4). Total number of variables obtained for each level is in parentheses. “Virulence, Disease and Defense” and its subsequent levels are shown as an example.

### Statistical analyses

Phylogenetic and functional abundances in percentages per sample were calculated using JMP Pro 11 (SAS Institute, Cary, NC). Discriminant analysis was used to evaluate changes in taxonomic composition (50 most abundant taxa) over time and by days after enrollment for each study group (*P-value* < 0.05). Stepwise selection was used and taxa with *P-value* < 0.1 were included in the discriminant model and in the subsequent multivariable mixed logistic regression model, described below.

Due to the large number of comparisons, the hierarchical data was screened for effects and only taxa or functions with a false discovery rate (FDR) *P-value* lower than 0.1 were selected for the multivariable mixed logistic regression model. For each category, a model was fitted for the most prevalent taxa or functions, the ones with significant FDR *P-values* or that were significant in the discriminant analysis. The independent variables study group (CON, ENR or TUL), sample day (4, 14, 56, and 112) and interactions were included as fixed effects in all models. Because values at enrollment diverged among the three groups, Day 0 measurements were included as a covariate in the model. The effects cohort and individual animals nested within cohort were controlled in the models as random effects. Figures were created using JMP Pro 11, GraphPad Prism 8.1.1 (GraphPad Software, San Diego, CA) and Microsoft PowerPoint 2016 (Microsoft Corporation, Redmond, WA).

## Results

### Shotgun sequencing

Whole genome sequencing of 179 samples produced 228,587,989 sequences totaling 57,397,337,159 basepairs (bp). The sample length was standardized to 251 bp when merging paired-end samples. One sample (calf in enrofloxacin group, Day 4) with a low number of sequences was excluded. Sequencing data by sample is listed in Supplementary Table S1 and more information is available at MG-RAST in project number 20043 (https://www.mg-rast.org/linkin.cgi?project=mgp20043).

### Effect of antibiotic treatment on taxonomic composition

Firmicutes, Bacteroidetes, Actinobacteria and Proteobacteria were the four major phyla and accounted for 96.4% of the microbiota on average. The top 20 most abundant phyla are presented in Figure 2 by sample day for each study group.

**Fig 2:**
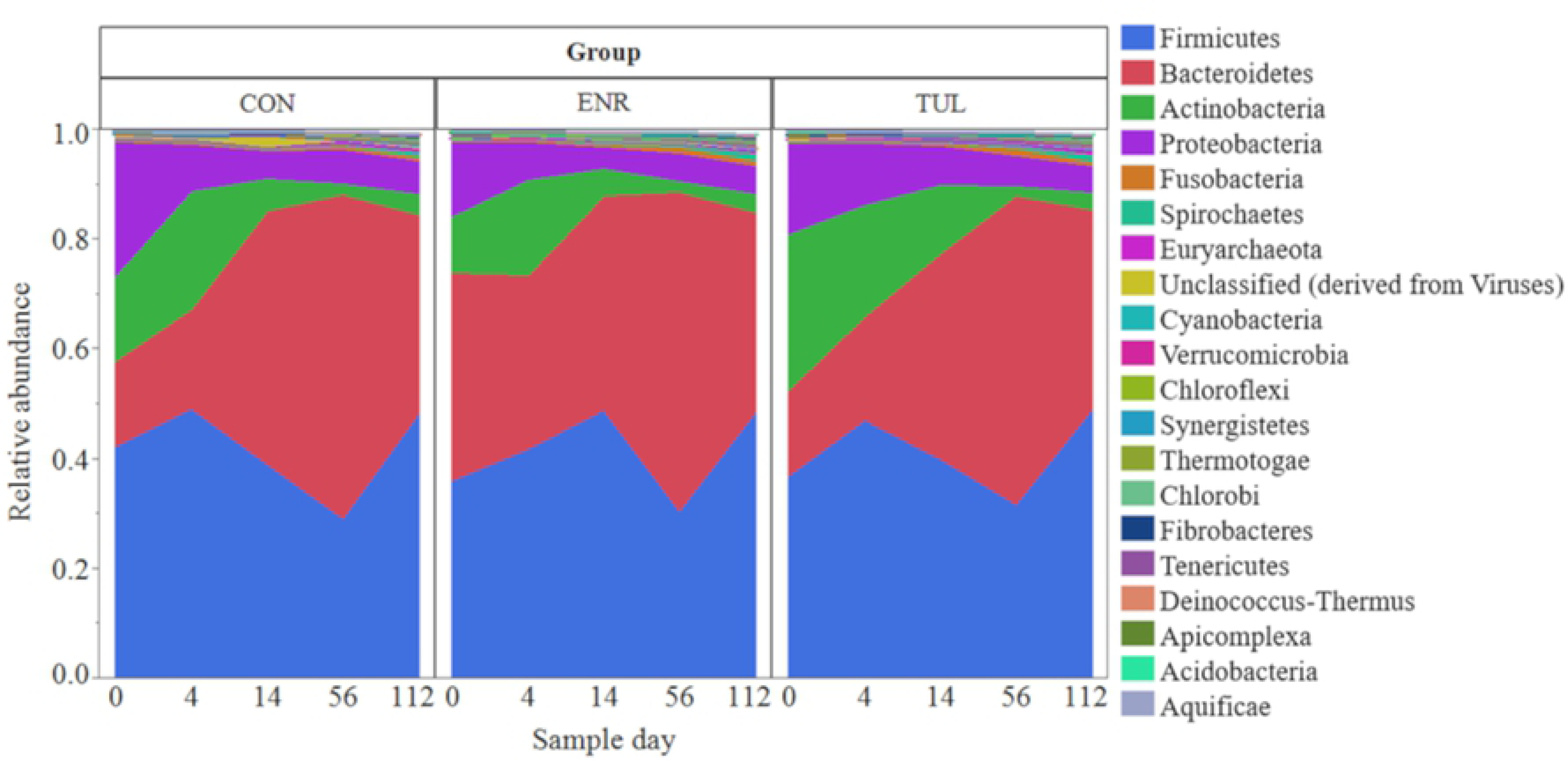
The top 20 most abundant phyla. The top 20 most abundant phyla by sample day for each study group (CON= control, ENR= enrofloxacin, TUL= tulathromycin).

Change in the microbiota composition was observed at the phylum level using discriminant analysis, as shown by the three-dimensional canonical plot (Figure 3). In this figure, each ellipse, which indicates the 95% confidence region to contain the true mean of the group of variables, represents a sampling day. The position of the ellipses shows how the microbiota changed over time, revealing a greater shift in microbial composition abundance observed from Day 14 to Day 112. When analyzing the same dataset by sample day, there was a difference at the phylum level on Days 0, with calves treated with enrofloxacin being more distant from the others, and on Day 112, when control calves were separated from the treated calves (Supplementary Figure S1). Discriminant analysis was performed at the subsequent phylogenetic levels. At the genus level, the differences were less pronounced compared to phylum level (Figure S2).

**Fig 3:**
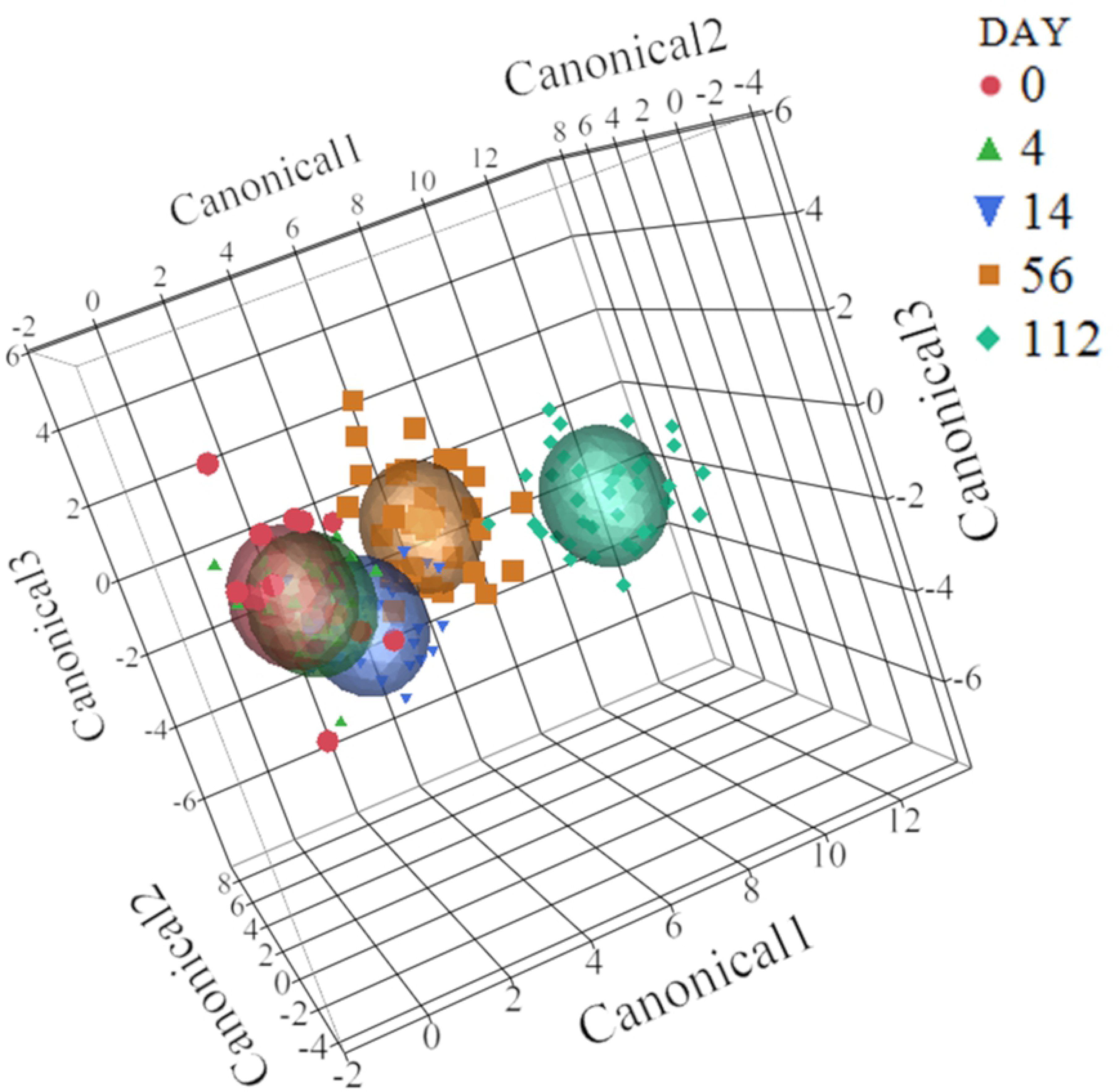
Discriminant analysis of the 50 most abundant phyla. Changes in phyla composition over time (*P-value* < 0.05). An ellipse indicates the 95% confidence region to contain the true mean of the variable (day).

When looking at the effect of antibiotic treatment on each taxon individually, including fixed and random effects, there was no significant effect of treatment group at the phylum level. As already described, because of the difference in baseline value between treatment groups on Day 0, it was included in the mixed model as a covariate.

At the class level, on Day 56 there was a numerical increase in the abundance of Deltaproteobacteria, which belongs to phylum Proteobacteria, for calves treated with enrofloxacin when compared to both other treatment groups (*P-value* 0.093). The abundance of Desulfovibrionales, an order within the Deltaproteobacteria phylum, was significantly different among groups, being higher on Day 56 in calves treated with enrofloxacin (*P-value* 0.015). The same significant trend was observed for the family *Desulfovibrionaceae* (*P-value* 0.010) and genus *Desulfovibrio* (*P-value* 0.009), as shown in Figure 4.

**Fig 4:**
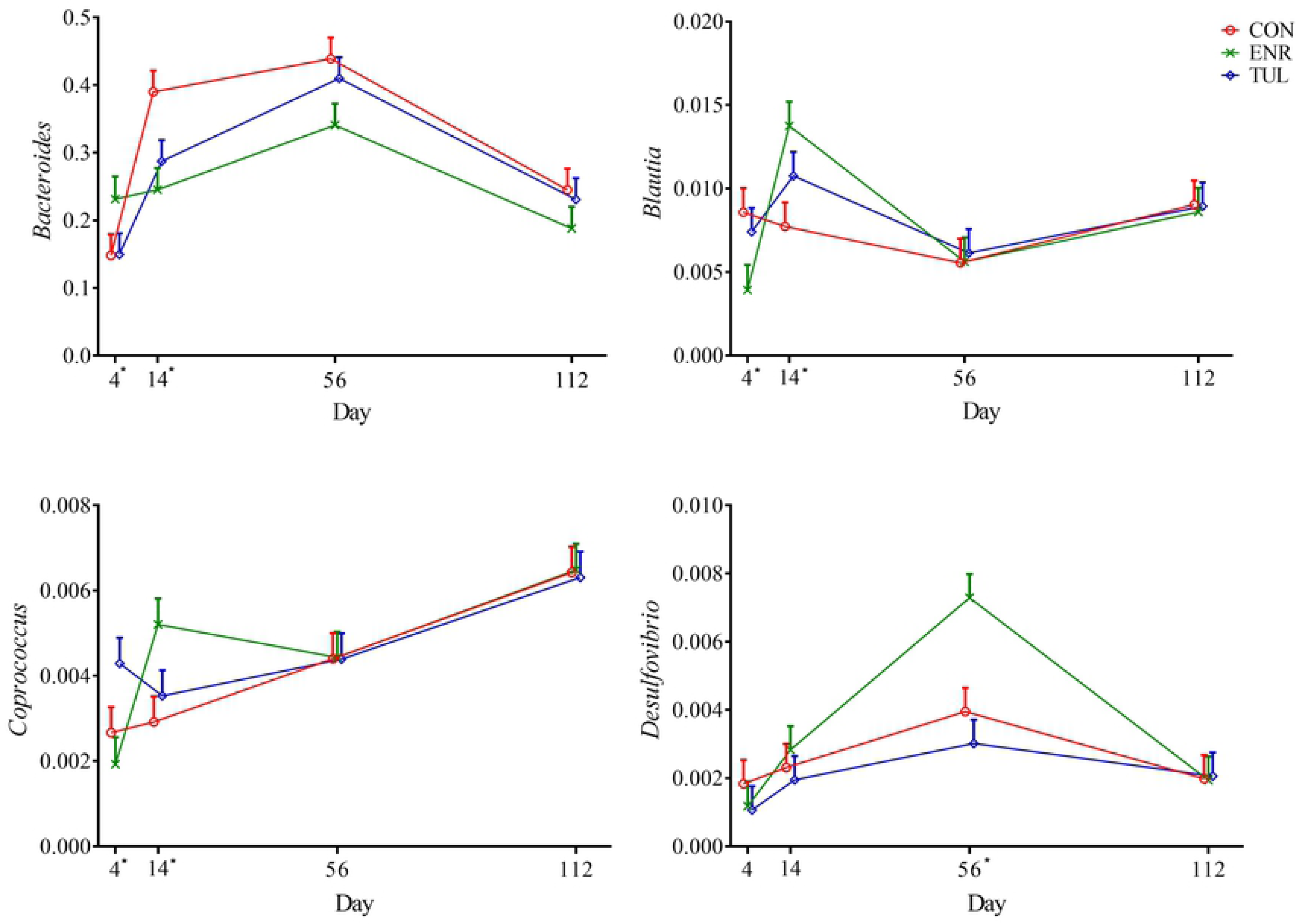
Relative abundance of genera *Bacteroides, Blautia, Coprococcus* and *Desulfovibrio* by sample day represented by least square means. The independent variables study group (CON= control, ENR= enrofloxacin, TUL= tulathromycin), sample day (4, 14, 56, and 112) and interactions were included as fixed effects in all models. Day 0 was included as a covariate in the model. The effects cohort and individual animals nested within cohort were controlled in the models as random effects. Asterisks indicate significant differences (*P-value* ≤ 0.05) for the sampling day. Error bars represent the standard error of the least square mean.

In the discriminant analysis, 16 unique genera were selected for the model (*P-value* < 0.1). When evaluated using the multivariable mixed logistic regression model, only *Bacteroides, Blautia, Coprococcus* and *Desulfovibrio* were significantly affected by treatment group and day (*P-values* 0.035, 0.019, 0.027 and 0.009, respectively; Figure 4).

### Effect of antibiotic treatment on microbiota function

As seen with microbial taxonomy, the discriminant analysis revealed a change in microbial function over time, which is shown by the three-dimensional canonical plot from Level 1 in Figure 5 (*P-value* < 0.05). As with microbial composition, the ellipses show a greater change in microbial function abundance as the animals became older.

**Fig 5:**
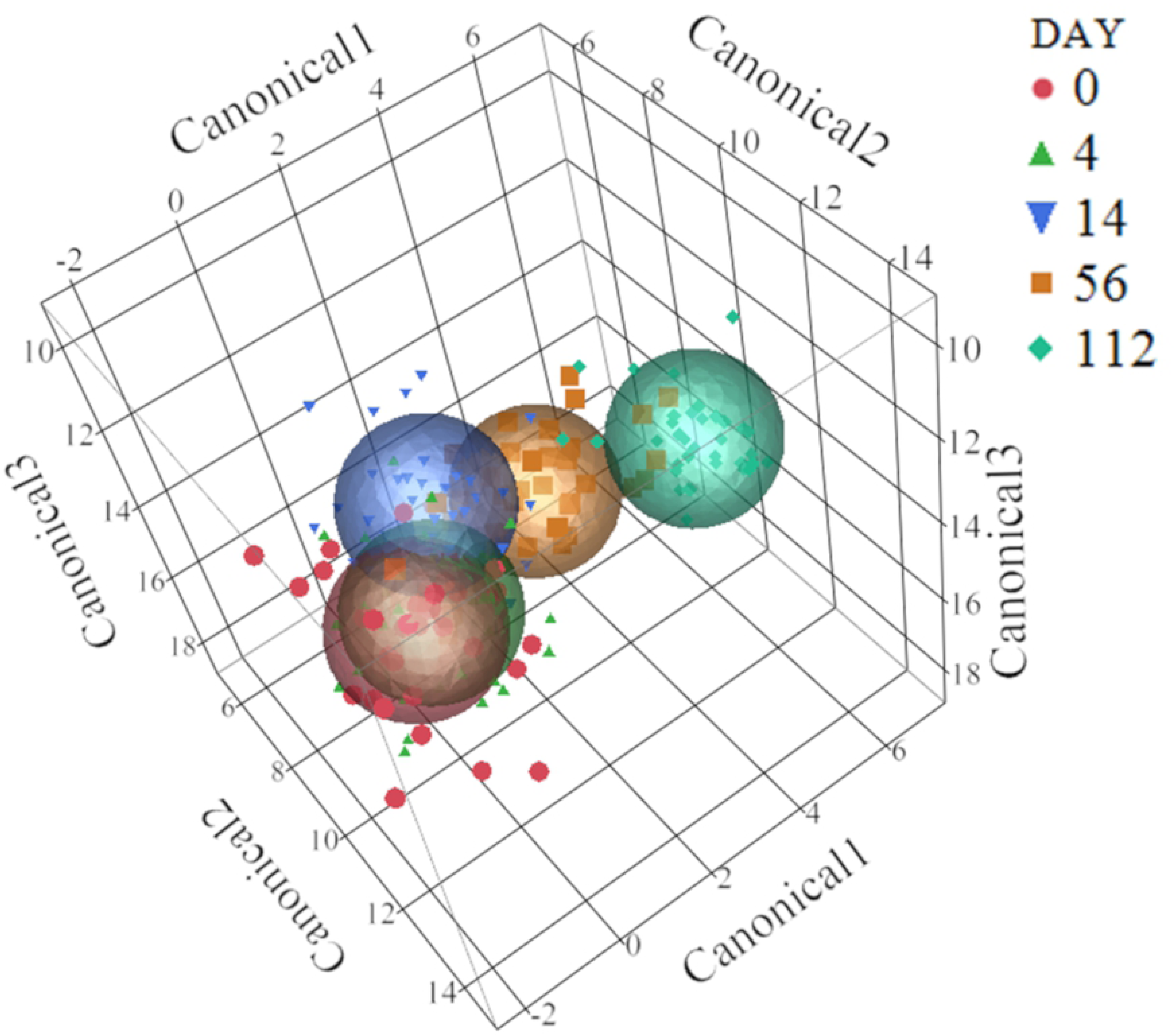
Discriminant analysis of the 28 microbial functions from Level 1. Changes in microbial function over time (*P-value* < 0.05). An ellipse indicates the 95% confidence region to contain the true mean of the variable (day).

At Level 1, the interaction between treatment group and day had a significant effect on the category “Virulence, Disease and Defense”; however, the difference was mainly on Day 0. When Day 0 was included in the model as a covariate, this significance was lost. Also at Level 1, there was a significant difference in the abundance of “Clustering-based subsystems” (*P-value* < 0.05), as shown in Figure 6. Despite differences observed over the first weeks, at Day 112 functions appeared to be at similar abundances among all study groups.

**Fig 6:**
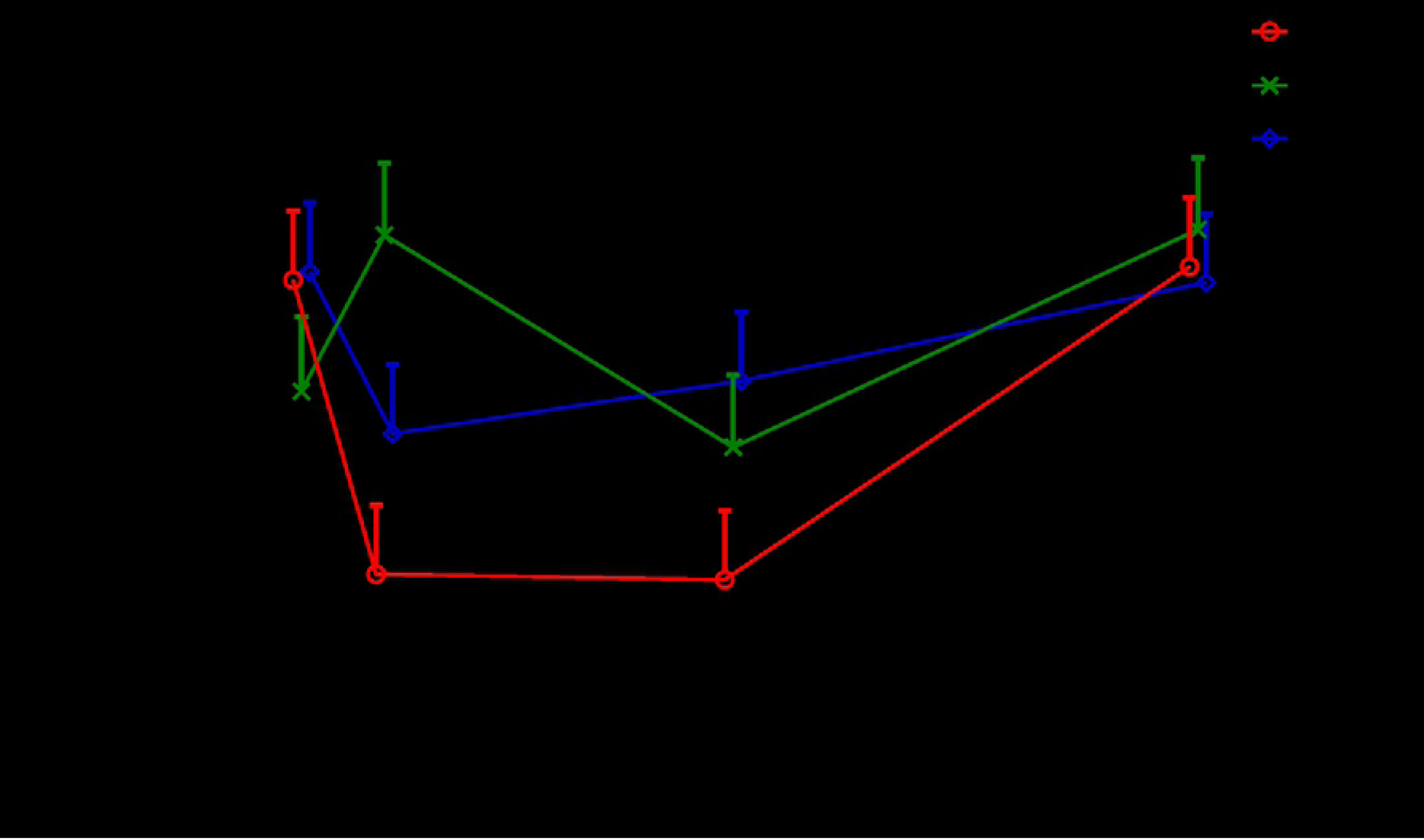
Relative abundance of genes with “Clustering-based subsystems” function by sample day represented by least square means. The independent variables study group (CON= control, ENR= enrofloxacin, TUL= tulathromycin), sample day (4, 14, 56, and 112) and interactions were included as fixed effects in all models. Day 0 was included as a covariate in the model. The effects cohort and individual animals nested within cohort were controlled in the models as random effects. Asterisks indicate significant differences (*P-value* ≤ 0.05) for the sampling day. Error bars represent the standard error of the least square mean.

At Level 2, calves in the enrofloxacin group had a higher abundance of genes with the “Transposable elements” function, which is part of Level 1 category “Phages, Prophages, Transposable elements, Plasmids”, on Day 4 (*P-value* 0.034) and then reached levels similar to the other two groups, as shown in Figure 7.

**Fig 7:**
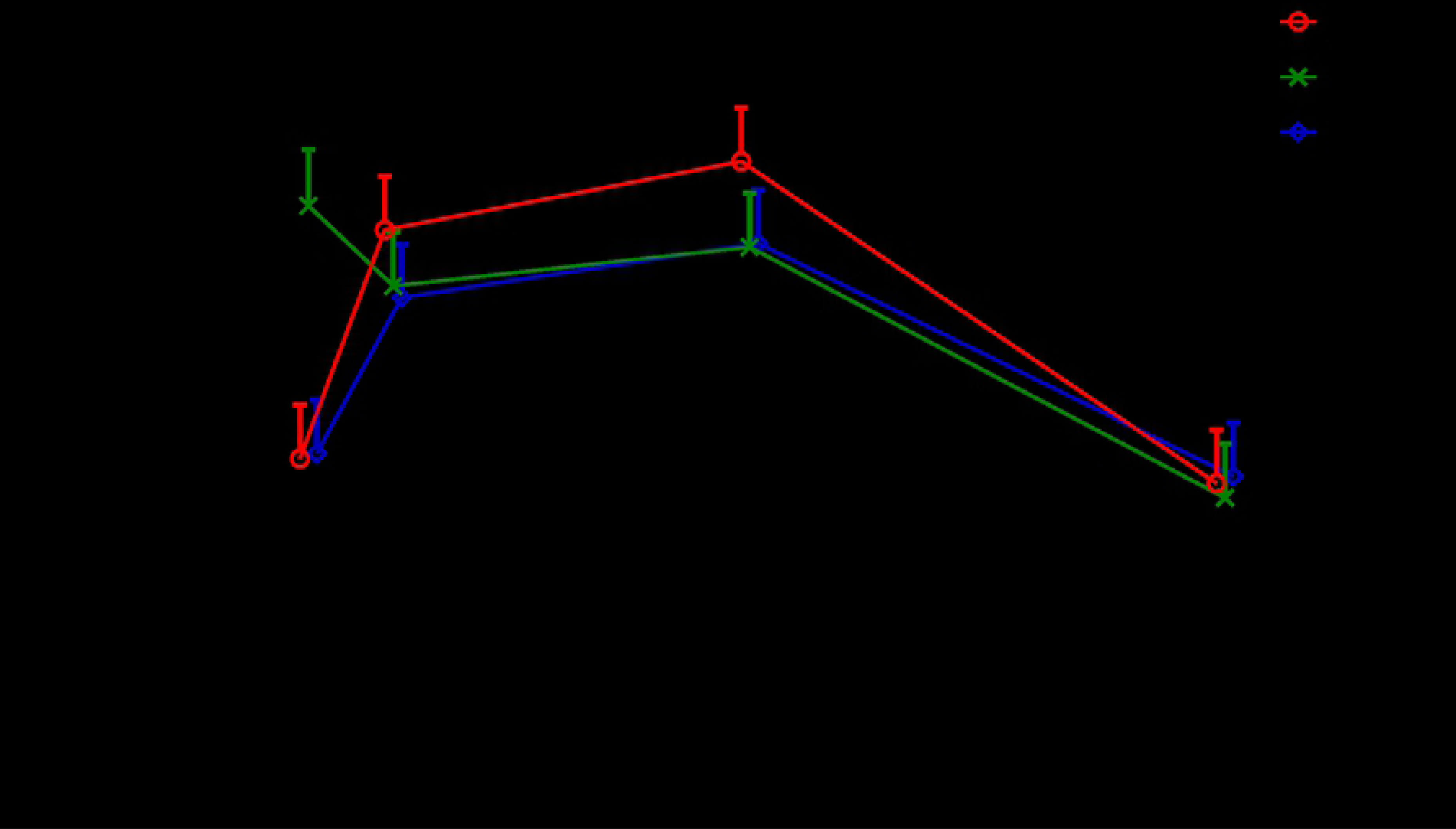
Relative abundance of genes with “Transposable elements” function by sample day represented by least square means. The independent variables study group (CON= control, ENR= enrofloxacin, TUL= tulathromycin), sample day (4, 14, 56, and 112) and interactions were included as fixed effects in all models. Day 0 was included as a covariate in the model. The effects cohort and individual animals nested within cohort were controlled in the models as random effects. Asterisks indicate significant differences (*P-value* ≤ 0.05) for the sampling day. Error bars represent the standard error of the least square mean.

The distribution of “Resistance to antibiotics and toxic compounds” (RATC) is depicted by the boxplots in Figure 8. There was a large variation in “RATC” abundance on Day 0, especially for the enrofloxacin group. Although calves that received enrofloxacin started with a higher abundance of “RATC” on Day 0, we did not observe a rise in AMR genes after treatment with enrofloxacin. According to the mixed model results, “RATC” was not statistically different among the three treatment groups (*P-value* 0.095). A graph and a table with the relative abundances and *P-values* can be found in Supplementary Figure 3.

**Fig 8:**
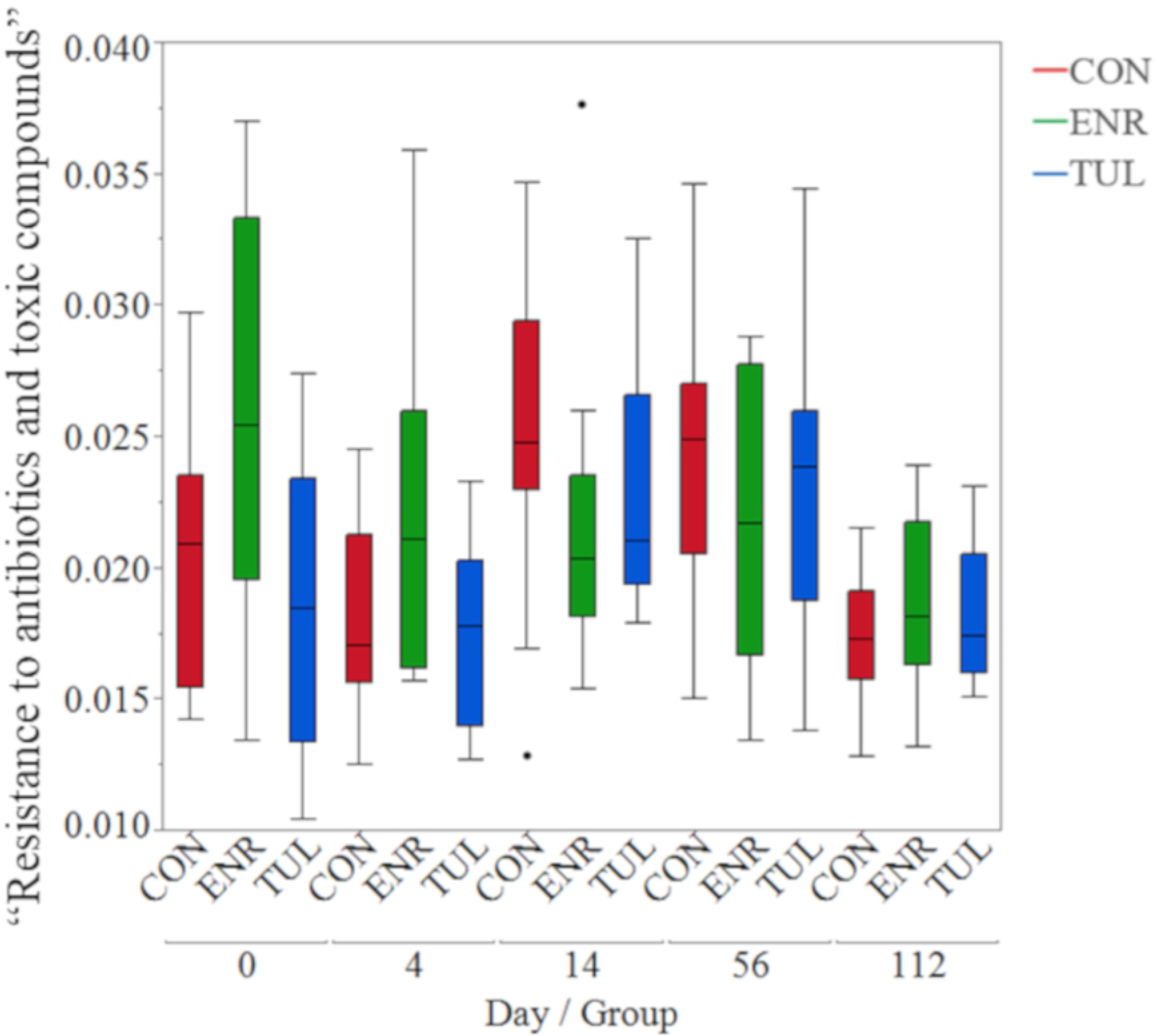
Boxplots of genes with “Resistance to antibiotics and toxic compounds” function. By sample day and study group (CON= control, ENR= enrofloxacin, TUL= tulathromycin).

At level 3, within “RATC”, we found no significant difference in the relative abundance of “Resistance to Fluoroquinolones” among treatment groups (*P-value* 0.19). Abundance of the “RATC” function increased over time for all treatment groups, including for calves that did not receive antibiotic treatment, and it was higher on Day 56 for calves treated with tulathromycin (Figure 9). Also within “RATC”, the relative abundance of “Multidrug efflux pump in Campylobacter jejuni (CmeABC operon)” was significantly different among groups (*P-value* 0.015), with calves treated with enrofloxacin having higher abundance on Day 4 and then lower on Day 56. There was no significant difference detected in “Erythromycin resistance” among treatment groups (*P-value* 0.662).

**Fig 9:**
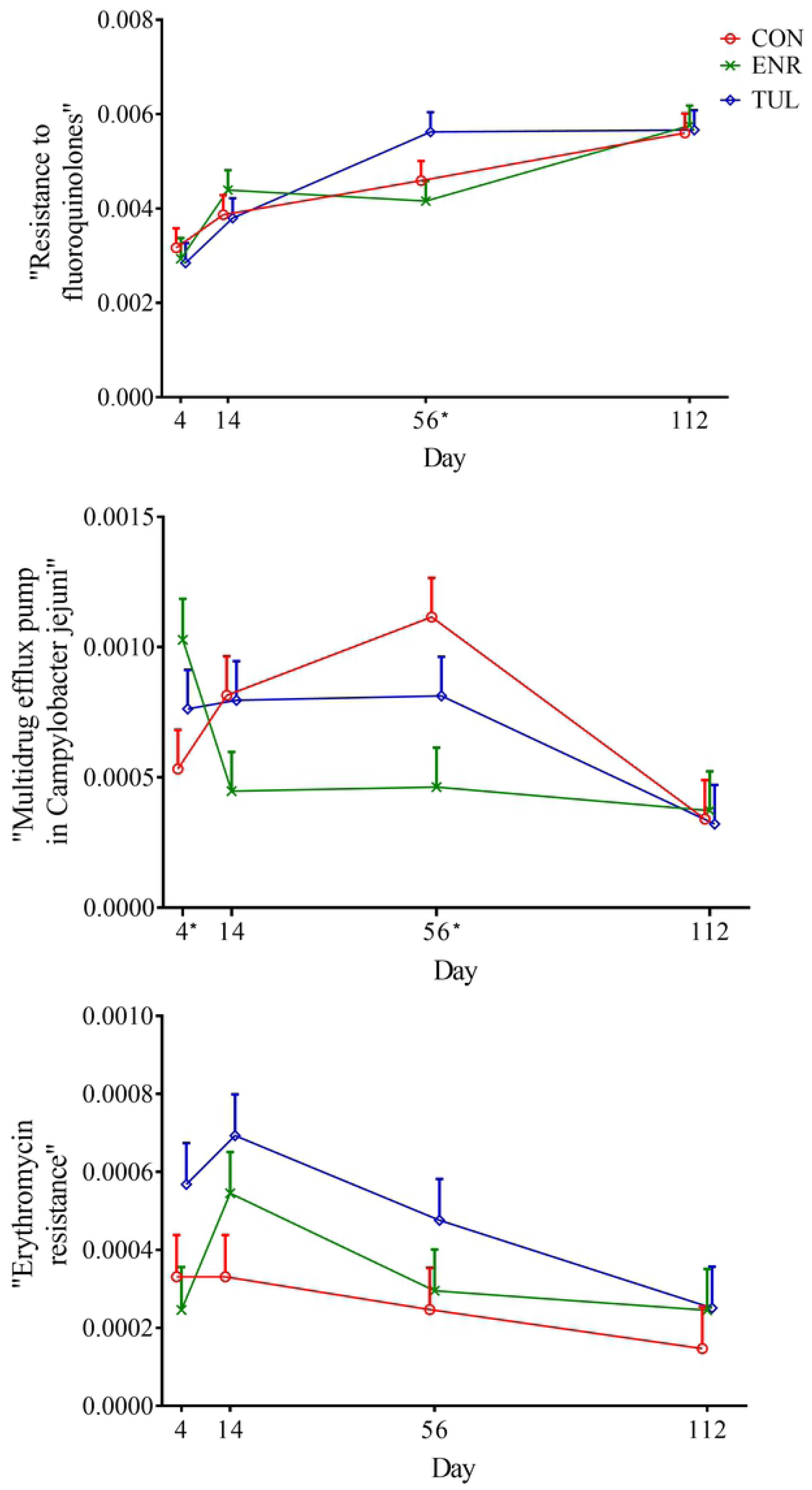
Relative abundance of functional categories within “RATC” by sample day represented by least square means. The independent variables study group (CON= control, ENR= enrofloxacin, TUL= tulathromycin), sample day (4, 14, 56, and 112) and interactions were included as fixed effects in all models. Day 0 was included as a covariate in the model. The effects cohort and individual animals nested within cohort were controlled in the models as random effects. Asterisks indicate significant differences (*P-value* ≤ 0.05) for the sampling day. Error bars represent the standard error of the least square mean.

At the fourth and lowest functional level, the relatives abundances of “DNA gyrase subunit A (EC 5.99.1.3)” within “Resistance to Fluoroquinolones” increased over time (*P-value* 0.037), and of “rRNA adenine N-6-methyltransferase (EC 2.1.1.48)” within “Erythromycin resistance” was significantly higher on Day 14 for the tulathromycin group, then it declined (*P-values* 0.026), as picture in Figure 10.

**Fig 10:**
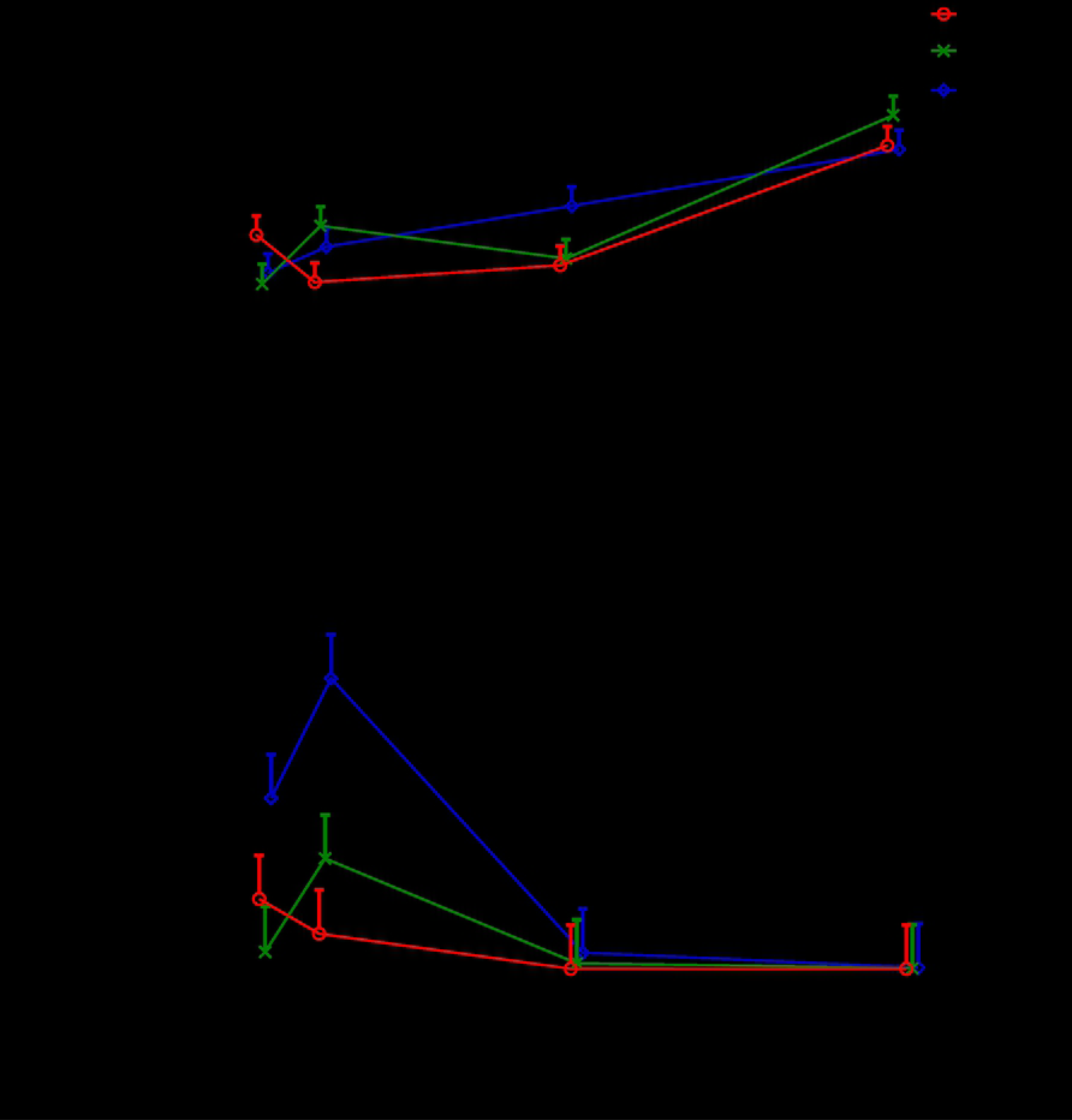
Relative abundance of functional categories at level 4, within “RATC”, by sample day represented by least square means. “DNA gyrase subunit A (EC 5.99.1.3)” and “rRNA adenine N-6-methyltransferase (EC 2.1.1.48)”. The independent variables study group (CON= control, ENR= enrofloxacin, TUL= tulathromycin), sample day (4, 14, 56, and 112) and interactions were included as fixed effects in all models. Day 0 was included as a covariate in the model. The effects cohort and individual animals nested within cohort were controlled in the models as random effects. Asterisks indicate significant differences (*P-value* ≤ 0.05) for the sampling day. Error bars represent the standard error of the least square mean.

## Discussion

Our results support that metaphylactic treatment of preweaned calves with enrofloxacin or tulathromycin did not cause major changes in fecal microbiota composition and gene functions related to antimicrobial resistance. Despite the fact that calves were not treated with antibiotics before enrollment and were randomly assigned to one of the three study groups, these groups started the trial with different microbiota composition and function, including higher abundances of “Virulence, Disease and Defense” and “RATC” functions in calves assigned to enrofloxacin treatment. Nevertheless, after controlling for this difference in the analysis, we did not observe a significant increase in the abundance of genes with these functions after treatment with antimicrobial drugs.

When lactating cows are treated with antimicrobial or other drugs due to illnesses, their milk is withheld from sale because it may contain drug residues. This non-saleable milk, also called waste milk, is often fed to calves. Antibiotic residues found in waste milk from 34 farms in New York state were mainly β-lactams, tetracycline and sulfamethazine [25]. Enrofloxacin is approved only for non-lactating female dairy cattle less than 20 months of age, and extra-label use of this drug is strictly prohibited; therefore, there should not be residues from this drug class in milk fed to calves. Additionally, enrofloxacin was not used at the study farm as a treatment for calf pneumonia before the study began.

Calves could have acquired drug resistant bacteria and resistant genes from their mothers during birth, from the waste milk used as feed [26,27] or from the environment [8,28], including cross-contamination through farm workers and feeding equipment [29]. Studies have found AMR genes in animals raised without antimicrobials as well [30,31]. Auffret *et al*. found differences in AMR genes depending on the diet (forage versus concentrate) fed to beef cattle not treated with antibiotics [31].

Treatment with antimicrobial drugs did not have a significant effect on “Virulence, Disease and Defense” and “RATC”, as we hypothesized. On the other hand, calves treated with enrofloxacin had higher abundance of “Transposable elements” on Day 4. Also known as transposons, these mobile genes can transfer functions within genomes and cause mutations [32]. The enrofloxacin group also had higher abundances of “Clustering-based subsystems” and lower abundances of “Multidrug efflux pump in Campylobacter jejuni (CmeABC operon)” on Day 14; however, it is not clear what these changes may represent.

More than 20 years ago, Brown *et al*. discussed the increase in resistance to fluoroquinolones over time and emphasized the need to use these drugs judiciously in human and veterinary medicine [33]. Cummings *et al*. alerted to an increase in enrofloxacin resistance in bovine *E.coli* isolates in the northeastern United States from 2004 to 2011 [34]. In a controlled trial evaluating effects of low concentrations of drug residues in waste milk, “Resistance to fluoroquinolones” was found on the first day of life and increased over the following 6 weeks, even in calves that did not receive any parenteral antimicrobial or drug residue in their milk [27]. In our study, “Resistance to fluoroquinolones” was present even before the use of enrofloxacin on Day 0, and increased over time, regardless of study group. This might be the result of co-resistance, when bacteria acquire resistance to more than one class of antibiotic [35].

Changes in taxonomic composition over time are expected as the animal grows, starts eating solids and the microbiome becomes more diverse. The age group of 2 to 3 weeks old was selected because of the higher risk for development of BRD at this stage of life, and enrofloxacin and tulathromycin are commonly used for treatment of heifers with BRD [11,36]. The parenteral use of these two antimicrobials did not cause significant deviations in the abundances of the four major phyla: Firmicutes, Bacteroidetes, Actinobacteria and Proteobacteria. These results are in agreement with other studies that evaluated fecal microbiome through shotgun and 16s rRNA sequencing [23,27,37,38], but in different proportions.

*Bacteroides* is a frequently-studied, gram-negative, anaerobic bacterial genus, composed of several species and strains including *B. fragilis, B. vulgatus, B. thetaiotaomicron*, and many others. [39,40]. They are abundant in the gut, but can be opportunistically pathogenic such as during trauma and post-surgical infections, [39,40]. *Bacteroides* can develop AMR to many antimicrobials, as extensively reviewed by Wexler *et al*. [41]. An increase in resistance to fluoroquinolones has been reported among *B. fragilis* group strains from humans in the United States (moxifloxacin) [42] and in Spain (moxifloxacin and trovafloxacin) [43]. Despite having more *Bacteroides* at Day 0, calves treated with enrofloxacin did not show a major increase in the relative abundance of genes related to “RATC”, while non-treated calves had an increase in *Bacteroides* abundance over time and in “RATC” as well.

Calves in the enrofloxacin group had an increase on Day 56 in the genus *Desulfovibrio*, its family and order. *Desulfovibrio* spp. are sulfate-reducing bacteria that produce hydrogen sulfide, which is both important for cell physiology and toxic to epithelial cells [44]. They are also opportunistic pathogens found in the environment [45] and GI tract of humans and other animals [46,47]. Because they are difficult to culture, there is a lack of information about this gram-negative anaerobe, including antimicrobial susceptibility data. In a study with 23 *Desulfovibrio* isolates from four different species cultured from human specimens, *D. fairfieldensis* strains showed resistance to β-lactams and the fluoroquinolones ciprofloxacin and levofloxacin [48]. Culture-independent studies will bring more knowledge to these pathogens, but culture-based methods continue to be important for antimicrobial susceptibility surveillance.

The genus *Blautia* has been associated with human gut health, being reduced in patients with colorectal cancer [49]. Together with other carbohydrate-utilizing bacteria, higher relative abundances of *Blautia* and *Coprococcus* were found by Song *et al*. in the gut of 3-week old dairy calves, which coincides with rumen development, solid intake and greater concentration of short-chain fatty acids [50]. These two genera were temporarily increased in calves that received enrofloxacin in our study, which may be considered a good, unpredicted side effect.

Pitfalls of our study include the fact that, although calves were randomly enrolled in treatment groups, initial microbial composition and function still differed between calves in different treatment groups at Day 0. This study was performed on a single farm, which did not allow comparisons of effects across herds. Nevertheless, the calf management practices of the farm used in the study are typical for many other commercial dairy farms in New York State.

In our study, enrofloxacin or tulathromycin had minimal impacts on the microbial composition and functional microbiota of calves over the study period (112 days) when used to prevent and control BRD. It is important to note that “Resistance to fluoroquinolones” increased during the study regardless of treatment group and, therefore, more efforts should be dedicated to reduce the use of medically important antimicrobials in dairy calves.

## Author Contributions

RVP, CA, LDW conceived and designed the experiments.

CF, RVP, JDS performed the experiments.

CF, RVP analyzed the data and wrote the paper.

All authors reviewed the manuscript. All authors read and approved the final manuscript.

### Acknowledgments

For their support and assistance, the authors thank Dr. Sabine Mann, Debora Pedroso, Laura M. Carroll, the Bicalho Lab, the Cornell University Biotechnology Resource Center – Genomics Facility and the Cornell Statistical Consulting Unit.

## Supporting Information

**Table S1:** Sequencing data by sample. More information is available at MG-RAST in project 20043 (https://www.mg-rast.org/linkin.cgi?project=mgp20043).

**Fig S1: Discriminant analysis of the 50 most abundant phyla**.

Changes for each study group on Day 0 and Day 112 after enrollment (*P-value* < 0.05). An ellipse indicates the 95% confidence region to contain the true mean of the variable (group). CON= control, ENR= enrofloxacin, TUL= tulathromycin.

**Fig S2: Discriminant analysis of the 50 most abundant genera**.

A) Changes in genera composition over time (*P-value* < 0.05). B) Changes for each study group by days after enrollment. An ellipse indicates the 95% confidence region to contain the true mean of the variable (day or group). CON= control, ENR= enrofloxacin, TUL= tulathromycin.

**Fig S3: Relative abundance of genes with “Resistance to antibiotics and toxic compounds” function by sample day represented by least square means**.

The independent variables study group (CON= control, ENR= enrofloxacin, TUL= tulathromycin), sample day (4, 14, 56, and 112) and interactions were included as fixed effects in all models. Day 0 was included as a covariate in the model. The effects cohort and individual animals nested within cohort were controlled in the models as random effects. Asterisks indicate significant differences (*P-value* ≤ 0.05) for the sampling day. Error bars represent the standard error of the least square mean.

